# Musical expertise enhances neural alignment-to-young in sensorimotor regions that predicts older adults’ audiovisual speech-in-noise perception

**DOI:** 10.1101/2022.11.05.515273

**Authors:** Lei Zhang, Xiuyi Wang, Yi Du

## Abstract

Musical training can offset age-related decline in speech-in-noise perception. However, how lifelong musical expertise affects the functional reorganization of older brain in speech-in-noise perception has not yet been systematically investigated. Here, we address this issue by analyzing fMRI responses of older musicians, older non-musicians and, young non-musicians identifying noise-masked audiovisual syllables. First, we confirmed that older musicians outperformed older non-musicians and even equaled young non-musicians. Then, we showed that both older groups showed decreased auditory activation and increased visual activation compared to young non-musicians, while older musicians showed higher activation in speech motor regions and greater deactivation of default mode network (DMN) regions than older non-musicians. Next, we revealed that musical expertise counteracted the age-related neural dedifferentiation of speech representation, making older musicians exhibit higher neural alignment-to-young in bilateral sensorimotor areas. Finally, we disentangled that greater activation in speech motor areas and stronger deactivation in DMN regions were correlated with higher neural alignment in sensorimotor areas, which strongly predicted better performance in older adults. Together, long-term musical expertise mitigates age-related deficits in audiovisual speech-in-noise processing through enhanced compensatory scaffolding that reserves youth-like representation in sensorimotor areas. Our findings provide a comprehensive perspective on understanding age- and experience-related brain reorganization during speech perception.

## Introduction

Among myriad cognitive deficits associated with aging, the difficulty of understanding speech in noisy environments is one of the major problems faced daily by the elderly, even if their peripheral hearing remains normal. Accumulating evidence suggests that age-related decline in speech-in-noise perception can be counteracted by long-term or short-term musical training^1–5^. However, we are still unclear about the mechanisms engendered by musical experience against the aging impact on speech perception.

According to the scaffolding theory of aging and cognition (STAC^6^), the adaptive brain responds to age-related structural and functional deterioration (e.g., shrinking cortical thickness, disrupted white matter integrity, neural dedifferentiation, for reviews, see Refs.^6–8^) by engaging compensatory scaffolding—the recruitment of additional circuitry that shores up declining structures whose functioning has become noisy, inefficient, or both. Life-course beneficial factors such as long-term musical training can ameliorate the negative impacts of structural and functional decline as well as enhance the compensatory scaffolding process^9^. Yet, the complex relationships between functional deterioration or reserve of original honed networks, engagement of additional scaffolding networks, and speech-in-noise perception performance have not been systematically investigated in older adults either with or without lifespan musical experience. This precludes a full understanding of the nature of age-related and musical experience-related brain reorganization in speech processing.

Older adults may adopt several potential mechanisms to cope with age-related auditory deficits. Firstly, they might recruit extra sensory and/or frontal regions to compensate. When processing audiovisual stimuli, older adults tend to maximize the use of multimodal information to compensate the declined unimodal perception (for reviews, see Refs.^10,11^). This strategy might be amplified by older musicians, due to their advantage in multisensory integration as a result of intensive multimodal (auditory, visual, motor) engagement in musical training. For instance, musicians show stronger auditory-motor integration during speech-in-noise perception^12^, stronger audiovisual integration (i.e., narrower temporal binding window) when processing flash-beep, music, and speech stimuli^13–15^, and more robust brainstem response to audiovisual speech^16^. Moreover, older adults recruit more frontal regions (e.g., inferior frontal gyrus, anterior insula, precentral gyrus) than young adults when perceiving speech in adverse conditions^17–20^, which is consistent with the posterior-anterior shift in aging (PASA) and decline-compensation hypothesis^21,22^. The greater recruitment of frontal areas might represent a compensatory scaffolding for greater listening effort, cognitive control, and sensorimotor integration. Besides, neural dedifferentiation, i.e., declined specificity of neural representations, is less observed in frontal speech motor regions (i.e., inferior frontal gyrus and ventral precentral gyrus) than in auditory areas in older adults during speech-in-noise perception. And older subjects with higher frontal activation show better preserve of speech specificity in both frontal and auditory regions, indicating that frontal up-regulation provides functional reserve for older adults^17^. Yet, it remains unknown whether over-recruitment of sensory and frontal regions provides sufficient compensation for performance in older musicians.

Secondly, although the age- and training-related changes in default mode network (DMN) regions and their role in speech-in-noise perception have rarely been investigated, we hypothesized that older adults need to inhibit DMN at a higher level than young adults. DMN is a set of widely distributed brain regions, showing reductions in activity during attention-demanding tasks but increasing activity during tasks linked to memory or abstract thought^23–25^. When performing the speech-in-noise task, older adults need to pay attention to external auditory information and inhibit task-unrelated long-term memory supported by DMN to avoid interference. The failure to inhibit DMN in the elderly has been shown to be detrimental to task performance^26,27^. Thus, it is likely that older adult who better inhibits DMN would exhibit better speech perception performance. While musical training could strengthen the brain’s executive function^28^, we would expect older musicians to deactivate DMN to a higher extent to reduce interference and allow attention allocation to the current task. Indeed, recent studies have found that musicians show strengthened structural and resting-state functional connectivity in DMN regions^29–31^.

The third potential mechanism is related to functional reserve, particularly for older musicians who may maintain youth-like brain activity patterns. It has been shown that musical training has a potential age-protecting effect on the brain. Compared to non-musicians, musicians had lower brain age scores, which means their predicted brain age is younger than their chronological age^32^. The youth-like brain activity pattern allows old musicians to functionally reserve auditory processing along the auditory pathway and cognitive abilities such as working memory(Alain et al., 2014; Bidelman and Alain, 2015; Dubinsky et al., 2019; Parbery-Clark et al., 2012; Zendel et al., 2019; Zhang et al., 2021). However, it remains an open but critical question whether and where in the brain reserved youth-like activity patterns contribute to maintained speech-in-noise performance in older adults, or whether the opposite patterns might be true, the less similar as young adults (i.e., stronger scaffolding of additional compensatory circuitry) the higher performance.

In this functional magnetic resonance imaging (fMRI) study, we aimed to uncover the age-related changes in brain activity patterns during audiovisual speech-in-noise perception and how lifespan musical experience influences the aging-related functional reorganization. Twenty-five older musicians (OM), 25 older non-musicians (ONM), and 24 young non-musicians (YNM) identified syllables in noise under three signal-to-noise ratios with congruent visual lip movements inside the scanner. As expected, ONM performed worse than YNM, but OM performed better than ONM and equally well as YNM. Then we tested the three above mechanisms. Firstly, we conducted univariate analysis and found that both older groups showed increased activation in visual areas and increased deactivation in DMN regions than YNM, while OM had stronger activity in frontal-parietal regions and greater DMN deactivation than ONM. Next, we used multivariate pattern analysis (MVPA) and neural alignment measurement defined by inter-subject spatial pattern correlation between each older individual and young brain average to examine neural dedifferentiation in ONM and youth-like brain activity patterns in OM. Results revealed degraded speech representations in bilateral sensorimotor areas in ONM compared to YNM, but reserved neural specificity of speech representations and high neural alignment-to-young in the same areas in OM. Lastly, for the first time, we found that higher activation in frontal speech motor areas and stronger deactivation in DMN supported the youth-like activity patterns in sensorimotor regions, which positively predicted behavioral performance in older adults. Our findings thus provide a strong linkage between different aspects of brain organization underlying aging and musical plasticity, highlighting youth-like activity patterns in bilateral sensorimotor regions as a core mechanism in supporting speech-in-noise perception in older adults.

## Results

### Musical expertise offset age-related decline in audiovisual speech-in-noise perception

As shown in Fig. 1, the mixed-design ANOVA revealed significant main effects of group (*F*(2, 71) = 22.89, *p* < 0.001) and signal-to-noise ratio (*F*(2, 71) = 133.94, *p* < 0.001), and their interaction (*F*(4, 71) = 7.10, *p* < 0.001) in perceiving speech-in-noise. Post hoc analysis showed that ONM had lower accuracy than YNM (t(71) = -6.62, p < 0.001, corrected), indicating that audiovisual speech-in-noise perception functions decline with age. But the decline trend was not seen in OM because OM and YNM had similar accuracy (t(71) = 2.18, p = 0.097, corrected) and OM had higher accuracy than ONM (t(71) = 4.48, p < 0.001, corrected). These results indicate that the age-related decline of audiovisual speech-in-noise perception could be relieved by lifelong musical training. However, no significant correlation was found between years or intensity (practice hours in a week) of musical training and behavioral performance (|r| < 0.28, Bayes Factor (BF) < 1.44).

**Fig. 1.**
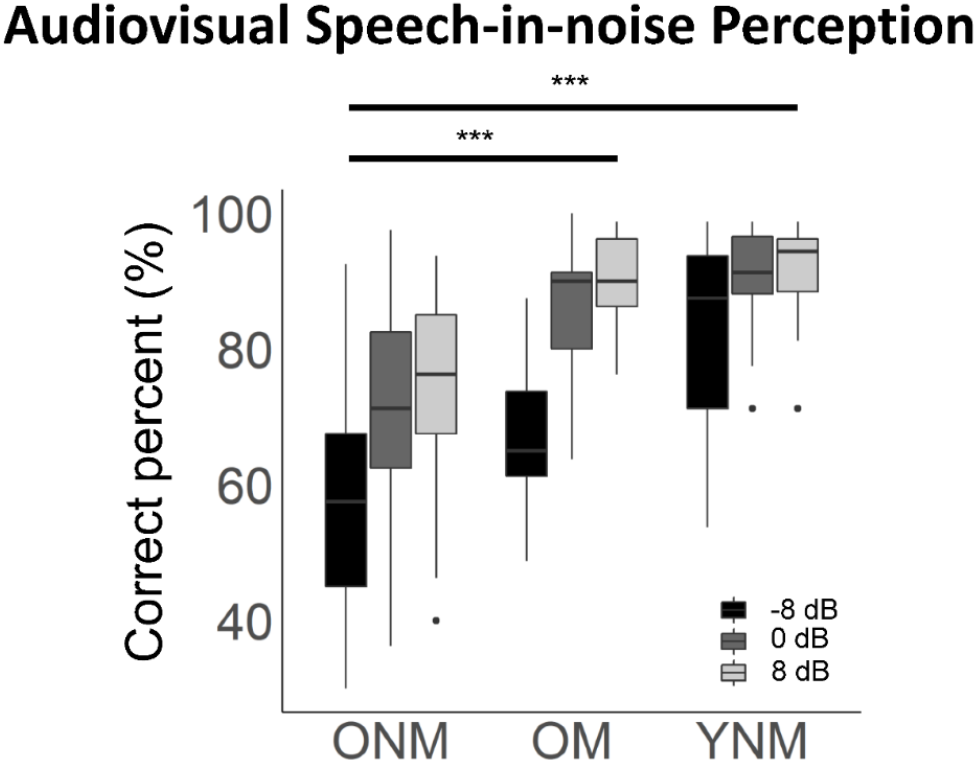
OM identified noise-masked audiovisual syllables equally well to YNM and better than ONM. Correct percent of audiovisual syllable-in-noise identification at 3 signal-to-noise ratios (-8, 0, 8 dB) in 3 groups. ONM, older non-musicians; OM, older musicians; YNM, young non-musicians. *** p < 0.001, by post hoc of mixed-design ANOVA, Bonferroni corrected.

### Older adults showed decreased auditory activities but increased visual activities and DMN deactivation during audiovisual speech-in-noise perception

To investigate the general aging effect on neural processing of audiovisual speech-in-noise perception, the task-related univariate blood oxygenation-level dependent (BOLD) activities were firstly compared between ONM and YNM, as well as between OM and YNM. Then, a conjunction analysis between OM vs. YNM and ONM vs. YNM was performed. No significant interaction between group and signal-to-noise ratio was found, and the effect of signal-to-noise ratio on neural activity was not reported in this study for simplifying the results and focusing on the group effects.

Both OM and ONM showed fewer activities in bilateral auditory areas but up-regulated activities in bilateral visual areas than YNM regardless of training experience (*p*_*fwe*_ < 0.05, Fig. 2a and Supplementary Table 1). Conjunction analysis confirmed that bilateral auditory and visual cortices were shared regions that exhibited a significant difference between older adults and young adults regardless of training experience (Supplementary Fig. 1 and Supplementary Table 2). However, auditory regions that showed decreased activation seemed wider in ONM than OM group.

**Table 1.** The group mean (standard deviation) values and statistics of age, education, pure tone average (PTA) at 250–4000 Hz, the Montreal Cognitive Assessment (MOCA) score, age of training onset, and years of music training in each group.

| Group | Age | Education | PTA | MOCA | Age of onset | Years of training |
| --- | --- | --- | --- | --- | --- | --- |
| Older musicians | 65.12<br>(4.06) | 13.02<br>(2.96) | 12.38<br>(5.39) | 27.92<br>(1.19) | 10.90<br>(4.56) | 50.88<br>(8.75) |
| Older non-musicians | 66.64<br>(3.40) | 9.50<br>(3.02) | 12.58<br>(3.58) | 27.52<br>(1.39) | NA | NA |
| t (p) | -1.43<br>(0.158) | 4.15<br>(<0.001) | -0.15<br>(0.881) | 1.09<br>(0.280) |  |  |
| Young non-musicians | 23.13<br>(2.38) | 16.50<br>(1.67) | 0.71<br>(3.32) | NA | NA | NA |

Note: Independent 2-sample t-tests were used for examining the group difference between older groups.

**Fig. 2.**
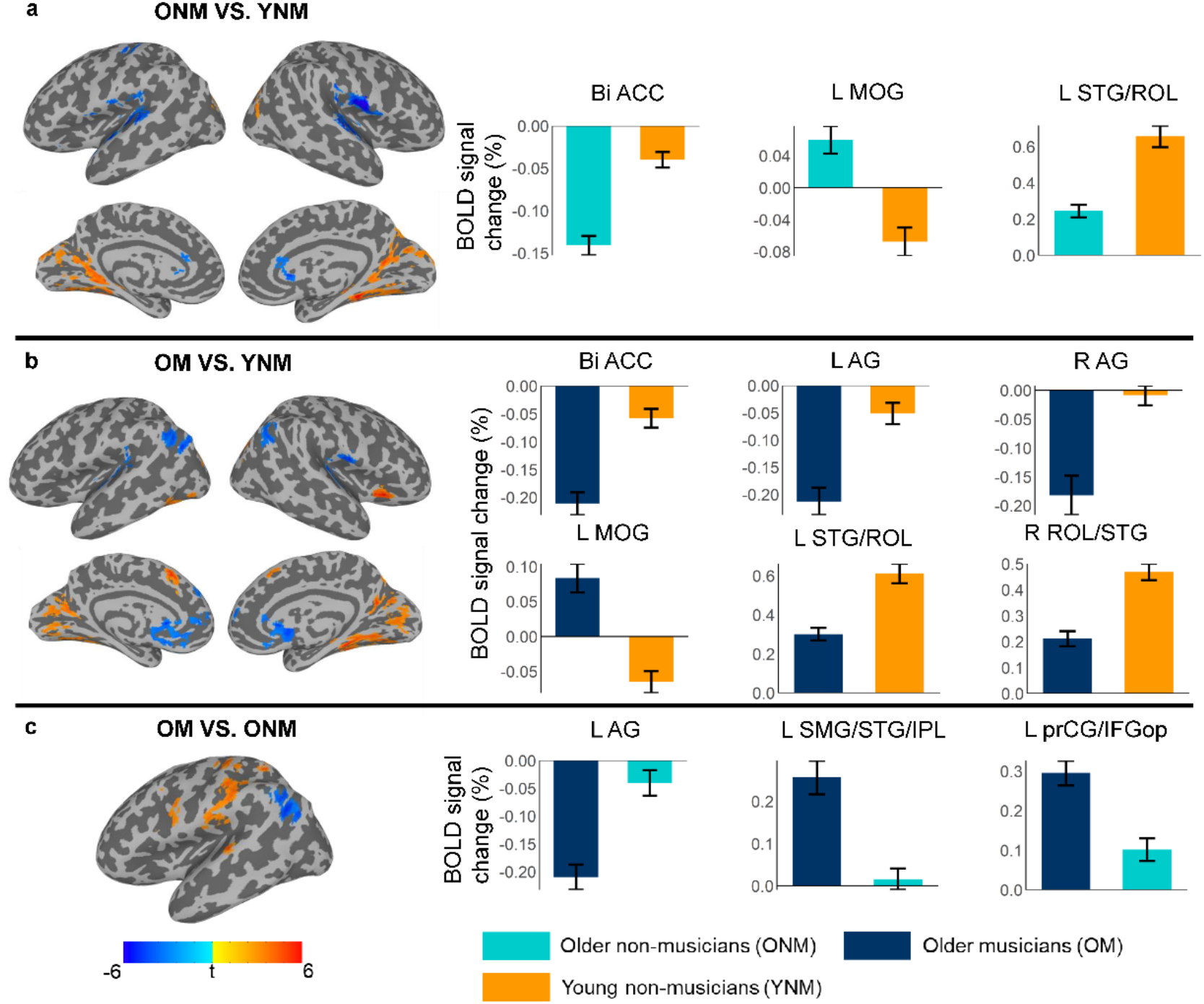
Group comparisons of BOLD activity (*p*_*fwe*_ < 0.05). (**a**) ONM showed greater activation in visual areas and less activation in auditory areas compared to YNM. (**b**) OM showed greater deactivation in DMN areas, greater activation in visual areas and less activation in auditory areas compared to YNM. (**c**) OM showed greater deactivation in DMN areas, and greater activation in frontal-parietal areas compared to ONM. Bar plots show the group mean BOLD signal changes in specific regions. Error bars indicate the standard error of the mean. Light blue: older musicians (OM), dark blue: older non-musicians (ONM), orange: young non-musicians (YNM). L, left; R, right; Bi, bilateral; ACC, anterior cingulate cortex; MOG, middle occipital gyrus; STG, superior temporal gyrus; ROL, rolandic operculum; AG, angular gyrus; SMG, supramarginal gyrus; IPL, inferior parietal lobule; prCG, precentral gyrus; IFGop, inferior frontal gyrus, opercular part.

ONM showed greater deactivation than YNM in bilateral anterior cingulate cortices (ACC), a core region of DMN^35^. OM showed greater deactivation in more DMN regions than YNM, including bilateral ACC, medial prefrontal cortex, and angular gyrus (*p*_*fwe*_ <0.05, Fig. 2a, b and Supplementary Table 1). Although both ONM and OM showed greater deactivation in bilateral ACC, conjunction analysis showed that OM deactivated more in the rostral part of ACC while ONM deactivated more in the caudal part of ACC (Supplementary Fig. 1 and Supplementary Table 2).

These results indicate that older adults might at least use two ways to compensate for declined auditory processing: (1) to fully utilize visual information and (2) to inhibit the task-unrelated regions, DMN, to reduce interference.

### Older musicians showed stronger frontal-parietal activation and DMN deactivation compared to older non-musicians

Having shown that OM performed equally well as YNM and better than ONM in audiovisual speech-in-noise perception, we investigated the counteraction mechanisms of lifelong musical training on age-related decline by comparing the activation between OM and ONM. Compared with ONM, OM showed stronger activation in left frontal-parietal regions, including the auditory-motor interfaces in posterior superior temporal gyrus and supramarginal gyrus, inferior and superior parietal lobule, and speech motor regions including the opercular part of inferior frontal gyrus (IFGop) of Broca’s area and the ventral premotor areas in precentral gyrus (prCG) (*p*_*fwe*_ <0.05, Fig. 2c and Supplementary Table 1). Musical training intensity of the first 3 years significantly predicted the activation of left IFGop (r = 0.42, BF = 6.21). Additionally, OM showed stronger deactivation in the left angular gyrus than ONM. Thus, long-term musical experience was associated with additional engagement of frontal-parietal regions and stronger deactivation of DMN regions in processing speech in noise.

### Neural representations of audiovisual syllables were degraded in older non-musicians but preserved in older musicians in visual and sensorimotor areas

How aging and musical experience altered neural representations of speech syllables were investigated using searchlight MVPA, which could examine the fine-scale spatial activation patterns rather than the overall neural activities during speech perception.

Although BOLD activities in visual areas were elevated in ONM compared to YNM, group comparison on MVPA classification accuracy showed that the neural specificity of phoneme representations still declined in ONM in left visual regions as well as in bilateral sensorimotor areas including prCG, postcentral gyrus, and Supplementary motor area (*p*_*fwe*_ <0.05, Fig. 3a and Supplementary Table 3). However, no significant result was found between OM and YNM, which means that the neural specificity of phoneme representations of OM was equivalent to that of YNM in visual and sensorimotor regions where speech representations were dedifferentiated in ONM. Indeed, directly compared with ONM, OM showed better audiovisual phoneme specificity in left visual areas and speech motor regions including prCG and IFGop (*p*_*fwe*_ <0.05, Fig. 3b and Supplementary Table 3).

**Fig. 3.**
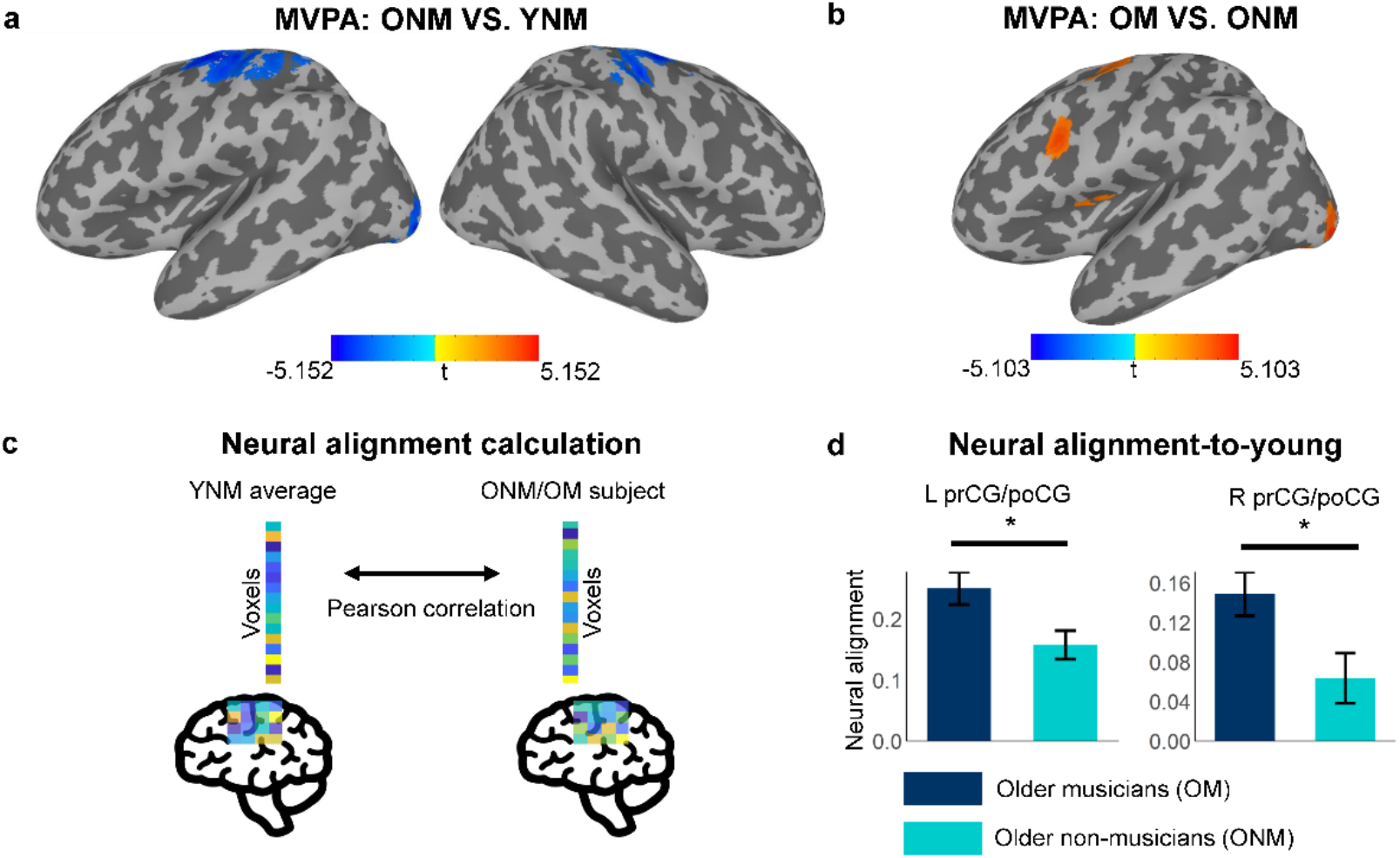
ONM showed degraded neural representation of audiovisual syllables, while OM kept similar neural specificity to YNM and higher neural alignment-to-young than ONM in sensorimotor areas. Clusters that showed significant difference in classification accuracy of syllables between ONM and YNM (**a**) and between OM and ONM (**b**). No classification difference was observed between OM and YNM. (**c**) Calculation of neural alignment-to-young. For each region with significant MVPA group difference, correlation between the mean spatial activation pattern of young adults and the pattern of each older adult was calculated for each syllable under each SNR and averaged across conditions to derive the neural alignment-to-young. (**d**) OM showed higher group mean neural alignment-to-young than ONM in left and right sensorimotor areas where significant group difference was found between ONM and YNM. Error bars indicate the standard error of the mean. * p < 0.05 by paired t-tests after correction. L, left; R, right; poCG, postcentral gyrus; prCG, precentral gyrus.

These results pinpoint the left visual areas and bilateral sensorimotor regions as the loci where age-related neural dedifferentiation of speech representation occurs and the left visual and speech motor areas as the target brain regions where musical experience-related offsetting of neural dedifferentiation presents.

### Older musicians showed higher spatial neural alignment to young brain than older non-musicians in bilateral sensorimotor areas

Since OM showed equivalent classification accuracy as YNM and both showed higher classification accuracy than ONM, we hypothesized that compared with ONM, OM would exhibit more similar activation patterns to YNM. To test this hypothesis, inter-subject spatial pattern correlation was further calculated between each older adult and the mean activation pattern of young adults for each audiovisual syllable inside the brain regions that showed significant group difference of classification accuracy (Fig. 3c, for details, see methods). This index provided a measure of the spatial neural alignment of multivoxel activation patterns of older brain to young brain template in the audiovisual speech-in-noise perception task.

As shown in Fig. 3d, OM showed significantly higher neural alignment-to-young than ONM in bilateral sensorimotor areas (significant regions in ONM vs. YNM group comparison of classification accuracy, left sensorimotor area: t(46) = 2.61, *p*_*fdr*_ = 0.039; right sensorimotor area: t(46) = 2.51, *p*_*fdr*_ = 0.039). Furthermore, musical training intensity of the last 3 years was positively correlated to neural alignment-to-young in bilateral sensorimotor areas (r = 0.39, BF = 4.46). Therefore, lifelong musical experience supports older adults to preserve higher level of youth-like neural patterns in bilateral sensorimotor regions during speech perception.

### Correlations among BOLD activities, neural alignment-to-young, and speech-in-noise perception performance

Our results suggest three compensatory mechanisms that older adults may adopt to process audiovisual speech in noise: 1) recruiting additional regions (i.e., frontal areas, visual regions), 2) inhibiting task-irrelevant DMN regions, and 3) maintaining similar activation patterns as young adults in sensorimotor regions. To further uncover which one was effective in compensation for behavioral performance and the relationships between those underlying mechanisms, correlation analyses were implemented pairwise between BOLD activities in sensory, DMN, and speech motor networks, neural alignment-to-young measurement, and speech-in-noise performance in two older groups. Since activations within the same neural network were highly correlated across older adults (Supplementary Fig. 2), principal component analysis was performed to extract activities in the correlated areas of each network into 1 principal component that was used for correlation analyses (for details, see Methods and Fig. 4a, b).

**Fig. 4.**
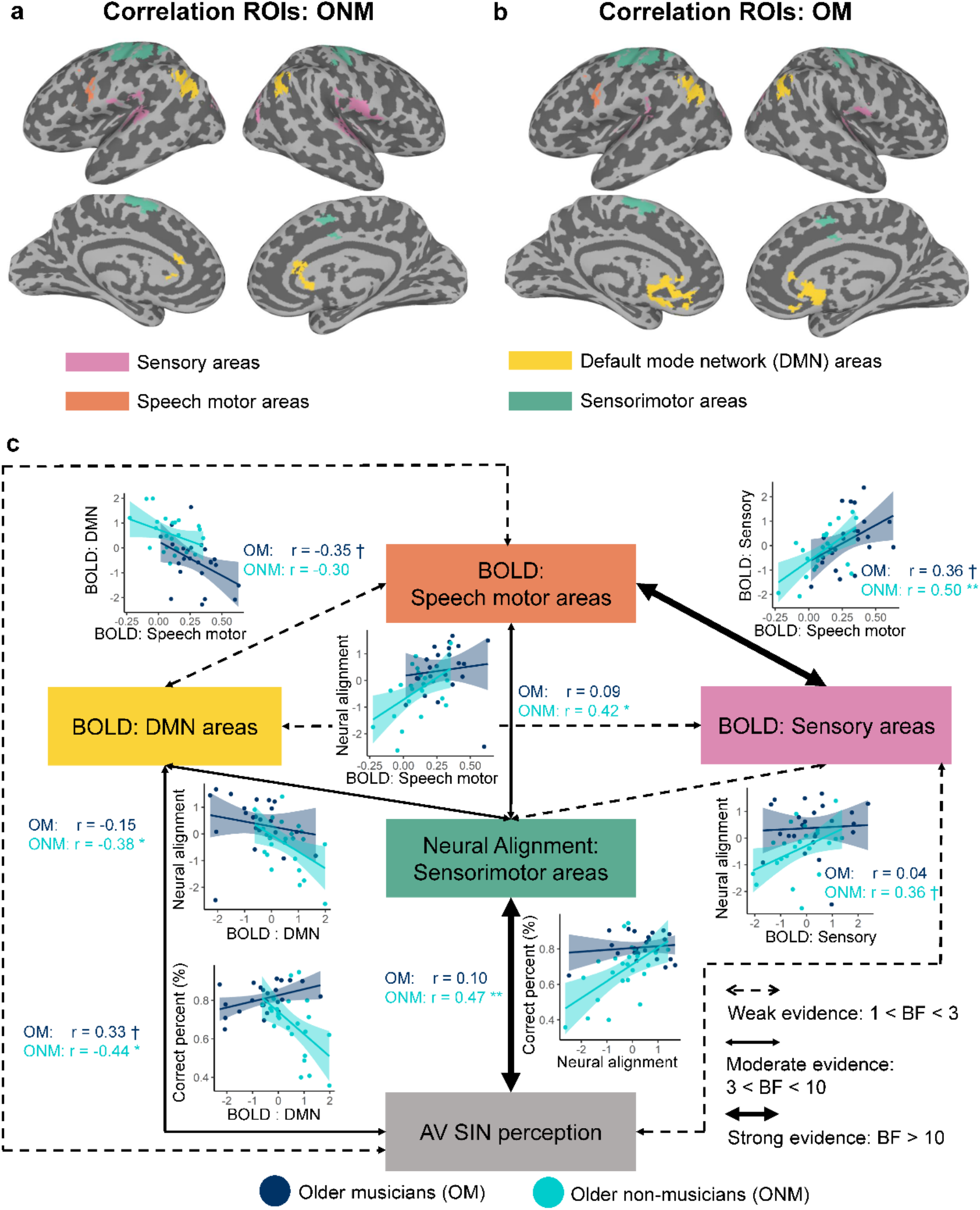
BOLD activation in speech motor areas and deactivation in DMN areas were positively correlated with neural alignment-to-young in sensorimotor areas, which positively predicted speech-in-noise performance in older adults. Regions where neural responses (BOLD activity in sensory areas, DMN areas, and speech motor areas, neural alignment-to-young measure in sensorimotor areas) were extracted in the correlation analysis for ONM (**a**) and ONM (**b**). Neural responses from regions with the same color were projected onto 1 principal component by principal component analysis if more than 1 cluster were included. (**c**) Scatter plots indicate Pearson correlation with Bayesian inference between 2 variables. Light blue: OM, dark blue: ONM. ** Bayes Factor (BF) > 10, * 3 < BF < 10, † 2.8 < BF < 3.

Regarding the first mechanism, additional engagement of visual and frontal speech motor regions did not compensate for speech-in-noise performance in the current study. As shown in Fig. 4c, BOLD activation in sensory areas (ONM: r = 0.08, BF = 0.48; OM: r = -0.30, BF = 1.74) and speech motor areas (ONM: r = 0.16, BF = 0.64; OM: r = -0.31, BF = 1.83) did not directly correlate with behavioral performance. However, activity in sensory areas positively correlated with activation in speech motor areas in ONM group (ONM: r = 0.50, BF = 21.30; OM: r = 0.36, BF = 2.76), which indicates the speech motor involvement as compensation for sensory engagement during speech-in-noise perception in older adults.

Regarding the second mechanism, greater DMN deactivation was correlated with better behavioral performance in ONM group but weakly correlated with worse behavior in OM group (ONM: r = -0.44, BF = 8.01; OM: r = 0.33, BF = 2.33). These results combining with the fact that OM exhibited stronger DMN deactivation than ONM (Fig. 2c) suggest that an intermediate level of deactivation in DMN areas supports the compensatory scaffolding of ONM for processing audiovisual speech-in-noise, but most OM exceed that level without extra behavioral benefit. Meanwhile, greater deactivation of DMN was weakly correlated with stronger activation in speech motor areas in OM group (ONM: r = -0.30, BF = 1.6; OM: r = -0.35, BF = 2.9), but not correlated with activity in sensory areas (ONM: r = 0.00, BF = 0.44; OM: r = -0.03, BF = 0.45), suggesting a synergistic relationship between DMN deactivation and speech motor engagement.

Regarding the third mechanism, the neural alignment-to-young in sensorimotor areas was found to strongly and positively predict audiovisual speech-in-noise performance in ONM group (ONM: r = 0.47, BF = 13.90, strong evidence; OM: r = 0.10, BF = 0.52), suggesting that higher youth-like brain activity patterns in bilateral sensorimotor areas helped older adults maintaining speech-in-noise performance. Meanwhile, stronger activities in speech motor areas and sensory areas, and stronger deactivation in DMN areas were correlated with higher neural alignment-to-young in sensorimotor areas in ONM group (speech motor: ONM: r = 0.42, BF = 5.51, moderate evidence, OM: r = 0.09, BF = 0.50; sensory: ONM: r = 0.36, BF = 2.99, weak evidence, OM: r = 0.04, BF = 0.61; DMN: ONM: r = -0.38, BF = 4.06, moderate evidence; OM: r = -0.15, BF = 0.61). Consistent with previous findings that neural alignment to the expert is highly correlated with learning performance^36^, the neural alignment to young brain in sensorimotor areas works as a hub linking functional preserve in sensory areas, compensatory scaffolding responses in speech motor and DMN regions to speech perception performance in older adults without musical experience.

## Discussion

This study investigated how aging and long-term musical training affect audiovisual speech-in-noise perception and the underlying neural preservation and reorganization. In line with and extending previous studies using audio-only speech stimuli(Alain et al., 2014; Dubinsky et al., 2019; Perron et al., 2022; Zendel and Alain, 2012; Zhang et al., 2021), we found that musical expertise offset age-related decline in audiovisual speech-in-noise perception. Paralleled with declined performance, older adults showed decreased activation in auditory areas, increased activation in visual areas, and stronger deactivation in DMN regions than young adults. In comparison, older musicians showed even greater deactivation in broad DMN areas than older non-musicians and young non-musicians. Older musicians also showed greater activation and better neural representation in frontal speech motor regions than older non-musicians. With neural alignment analysis, we are the first to show that the representation patterns in bilateral sensorimotor areas of older musicians was more similar to those of young adults compared to older non-musicians. Most importantly, stronger activation in speech motor areas and deactivation in DMN areas predicted higher neural alignment to young brain in sensorimotor regions, which was correlated with better behavioral performance in older non-musicians. Our findings thus pinpoint youth-like representation patterns in bilateral sensorimotor areas as a core mechanism of lifelong musical experience-related plasticity, linking retained sensory activity, compensatory scaffolding of the frontal speech motor and DMN regions with speech-in-noise perception performance in older adults. By tying up all the loose ends, this study provides a comprehensive understanding of age-related decline and compensation in speech processing and how musical expertise counteracts the effects of aging.

According to the STAC model, neural resources are depleted with aging, while compensatory scaffolding provides additional protection against cognitive decline^9^. In this study, compared with young adults, both older groups showed decreased activation in auditory areas during audiovisual speech-in-noise perception, indicating neural resource depletion with aging. However, both groups showed increased activation in visual areas, suggesting that they might recruit visual areas for compensation. Recruiting visual information is potentially beneficial in the current audiovisual speech-in-noise perception task in which both the auditory and visual information might provide important clues about the phoneme. Older adults are more susceptible to multisensory illusions, such as the sound-induced flash illusion and the McGurk illusion, suggesting that they are more inclined to use multimodal information and show stronger multimodal integration than young adults (for reviews, see Refs.^10,11^). During audiovisual speech perception, the contribution of auditory and visual information is weighted according to their reliability^37^. The decrease in auditory activation and increase in visual activation may underlie older adults’ tendency to utilize visual information to compensate for degraded auditory information. However, activation in sensory areas did not correlate with behavior performance in both older groups. Moreover, as revealed by MVPA analysis, the neural specificity of audiovisual speech in visual areas declined in older non-musicians compared to older musicians and young non-musicians. These results indicate that although older adults give greater weight to reliable visual information, only older musicians could maintain the specificity of neural representation of audiovisual speech in visual areas, and changes in sensory engagement cannot provide sufficient compensation for speech-in-noise performance.

In addition, for the first time, we found that older adults exhibited a greater deactivation in wide DMN areas including bilateral ACC, medial prefrontal cortex, and angular gyrus during the task than young adults. The DMN regions, which are involved in internal cognitive processes related to memory and abstract thought, are found to be typically deactivated when subjects perform goal-directed tasks^23^, with an anticorrelation between DMN activity and task performance^38^. The deactivation in DMN reflects an active suppression of irrelevant information processing when the environment demands externally focused attention rather than a relative increase in metabolic activity during rest^39^. DMN deactivation has been found to support attentional reallocation to external stimuli, which is critical for goal-directed activities^24,40^. Here, the higher deactivation level in DMN regions was correlated with better performance for older non-musicians, suggesting that stronger DMN deactivation helps older adults to better concentrate on the external speech stimuli and inhibit task-unrelated long-term memory supported by DMN to avoid interference. However, we found a weak negative relationship between DMN deactivation and behavioral performance in older musicians. Considering that the overall deactivation level of DMN regions was relatively high in older musicians, we speculated that a moderate level of DMN deactivation was optimal for reallocating attentional resources to external speech stimuli to achieve the best performance in older musicians. Overall, stronger DMN deactivation during audiovisual speech processing is a compensatory scaffolding strategy against aging and enhanced by lifelong musical experience.

Besides, compared to older non-musicians, older musicians exhibited higher activation in frontal-parietal regions belonging to the auditory and visual dorsal stream and better speech representation in speech motor areas. The visual dorsal stream is regarded as a “vision-for-action” stream that mediates motor programming and control based on current visual information^41,42^; while the auditory dorsal stream is considered a “sound-to-action” stream that maps auditory features to motor articulatory sequences^43^. Speech perception under adverse conditions particularly recruits speech motor areas in the dorsal stream including premotor cortex in prCG and IFG to compensate for deteriorated auditory processing via sensorimotor integration^44^. With visual lip movements, better speech encoding and tightened functional connectivity along the auditory and visual dorsal stream contribute to enhanced speech-in-noise perception^45,46^. Since musical training requires intensive interaction between sensory and motor modalities, musicians are found to exhibit stronger sensorimotor integration than non-musicians during perception^12,47–52^. Therefore, the enhanced recruitment of auditory and visual dorsal stream regions and better neural representation of audiovisual speech in speech motor areas by lifelong musical experience indicate that older musicians could better employ internal motor plans and external visual lip movements to aid speech-in-noise perception through strengthened visual/sound-to-action mapping and sensorimotor integration. However, mere activity in speech motor areas was not correlated with behavioral performance in either of the older groups, suggesting that there is an additional strategy that directly supports performance in older adults.

It has been shown that amateur musicians have younger brains compared to non-musicians^32^ and novices who show better neural alignment to experts have better learning outcomes in adults and children^36,53^. As expected, compared with young non-musicians, older non-musicians showed decreased neural specificity of phoneme representation, (i.e., neural dedifferentiation, see Refs.^7,8^), in bilateral sensorimotor regions. However, in the same regions, older musicians showed similar neural specificity of phonemes to young non-musicians and older musicians showed higher neural alignment-to-young than older non-musicians. Motor and somatosensory cortex are not only involved in speech production but also involved in speech perception^54,55^. Several neuroimaging studies have revealed that motor and somatosensory cortex represent phonemic features that facilitate speech perception in adverse listening conditions^44,54,56^. Therefore, our results suggest that musical expertise can reduce the age-related neural dedifferentiation and maintain youth-like activity patterns in sensorimotor areas, which is assumed to be critical for speech perception in noise. Importantly, we also found that greater neural alignment to young adults in sensorimotor regions was strongly correlated to better performance in older non-musicians, supporting that older adults may rely more on functional reserve of key brain regions in maintaining speech perception function. These results argue against the functional compensation hypothesis that lower neural alignment-to-young gives rise to better performance in older adults. By contrast, our results support the functional reserve hypothesis that higher neural alignment-to-young gives rise to better performance in older adults.

Furthermore, rather than demonstrating multiple neural changes associated with aging and musical experience individually, this study merited a panoramic illustration of the relationships among those neural alterations and behavior. We found that stronger activation in speech motor areas and sensory areas (weak effect) and stronger deactivation in DMN regions were positively correlated with better alignment-to-young in older non-musicians. Thus, the retained activation of sensory cortices, the additional engagement of frontal speech motor areas, and the inhibition of DMN regions converged on the maintaining of neural alignment-to-young in sensorimotor areas, which functioned as a critical predictor of audiovisual speech-in-noise performance in older adults. We failed to find any significant correlation between the BOLD activity and the alignment-to-young or between the alignment and behavior in older musicians. This may be due to the small variance of the alignment index in the older musician group. Still, lifespan musical experience enhanced compensatory scaffolding of older adults which supported their youth-like neural activation pattern, and the high alignment-to-young level resulted in their equivalent audiovisual speech-in-noise perception performance.

A few limitations exist in the current study. Firstly, only local activation and speech representation were analyzed here. To further unravel the age- and experience-related functional alterations of the neural networks during audiovisual speech perception, the group difference in functional connectivity needs to be investigated in further studies. Secondly, since older musicians who were eligible and willing to participate in our fMRI study are rare, older musicians with different types of musical training were combined into 1 group (e.g., piano, singing, violin, etc.). However, comparing the effect of different types of lifelong musical training that older adults majored in could be a valuable question to further reveal how musical training promotes multisensory speech perception more specifically because different training types require different kinds of sensorimotor integration (e.g., singing engages articulation system, playing piano engages fingers, etc.).

In summary, this study provides an integrative view in understanding the age-related decline-compensation in perceiving audiovisual speech in noisy circumstances and how lifespan musical experience offsets age-related decline by enhancing compensatory scaffolding and preserving youth-like brain. Multiple task-related functional changes were revealed with aging, including deficient recruitment of auditory areas but increased engagement of visual regions, enhanced DMN deactivation, and neural dedifferentiation of speech representation in sensorimotor and visual regions. In contrast, lifelong musical expertise was associated with further strengthened DMN deactivation, additional recruitment of speech motor areas, and maintained higher youth-like representation patterns in bilateral sensorimotor regions. Most importantly, the neural alignment-to-young in sensorimotor areas served as a key mediator by both aging and musical expertise, bridging the adaptive activation levels in sensory, speech motor, and DMN regions with behavioral performance in older adults, with more reserved youth-like patterns contributing to better performance. Our findings provide new insights into brain reorganization in the aging population and how long-term training experience affects it.

## Methods

### Participants

Twenty-five OM (65.12 ± 4.06 years old, 11 females), 25 ONM (66.64 ± 3.40 years old, 16 females), and 24 YNM (23.13 ± 2.38 years old, 12 females) completed this study. One OM with excessive head motion (more than 50% data was censored in the 3ddeconvolve) and 1 ONM who was left-handed were excluded from the fMRI analysis. The rest of participants were all healthy, right-handed, native Chinese speakers with no history of neurological disorder and normal hearing (average pure-tone threshold < 20 dB HL from 250 to 4,000 Hz) in both ears. OM were recruited from the conservatory of music, chorus, and orchestras. All participants had signed the written consent approved by the Institute of Psychology, Chinese Academy of Sciences. OM had started training before 23 years old (mean = 10.90 ± 4.56 years old), had at least 32 years of training (mean = 50.88 ± 8.75 years), and practiced consistently in recent 3 years (1 to 42 hours per week, mean = 12.70 ± 8.99 hours per week). Non-musicians reported less than 2 years of musical training experience. All older adults passed the Montreal Cognitive Assessment (MOCA) of Beijing version (≥ 26 scores) ^57^. Age, pure tone average, and MOCA score were balanced between OM and ONM (see Table 1). However, OM showed more years of education than ONM (t = 4.15, p < 0.001). Note that years of education of older adults here included informal education like on-the-job training. We conducted a few Supplementary analyses to exclude the effect of education on our results (for details, see section 1 of the Supplementary materials).

### Experimental design

The audiovisual stimuli comprised 4 naturally pronounced consonant-vowel syllables (ba, da, pa, ta) uttered by a young Chinese female. The utterances were videoed in a soundproof room. Videos of each syllable lasted 1s and were digitized at 29.97 frames/s in 1024 × 768 pixels. The pictures of the videos were cut retaining the mouth and the neck part. The audios were low-pass filtered (4 kHz), matched for average root-mean-square sound pressure level, and aligned in time by lip movement onset (started around the 10th frame of the video). The audiovisual syllables were masked by a speech spectrum-shaped noise (4 kHz low-pass, 10ms rise–decay envelope) that represents the spectrum of 113 different sentences by 50 Chinese young female speakers at 3 signal-to-noise ratios: -8, 0, and 8 dB (for details, see Refs.^45^). In the fMRI scanner, subjects were instructed to listen to the speech signals, watch the mouth on the screen, and identify the syllables by pressing the corresponding button using their right-hand fingers (index to little fingers in response to ba, da, pa, and ta in half of the subjects, pa, ta, ba and da in the other half of the subjects sequentially). Each subject completed 4 blocks. Each block contained 60 stimuli (20 trials × 3 signal-to-noise ratios), which were pseudo-randomly presented with an average inter-stimuli-interval of 5 s (4–6 s, 0.5 s step). Stimuli were presented via Psychtoolbox^58^.

### Behavioral analysis

To examine the group difference in behavioral performance, a 3 by 3 mixed-design ANOVA was performed (within-subject variable: signal-to-noise ratios; between-subject variable: subject group). Post hoc analyses were performed to further investigate the differences between the 2 groups. The Bonferroni correction was applied to correct the multiple comparisons. Statistical analysis was conducted in R^59^ with the package bruceR^60^ and visualized using the package ggplot2^61^.

### Functional imaging data acquisition and preprocessing

Imaging data were collected by a 3T MRI system (Siemens Magnetom Trio). T1 weighted images were acquired using the MPRAGE sequence (TR = 2200 ms, TE = 3.49 ms, FOV = 256 mm, voxel size = 1×1×1 mm). T2 weighted images were acquired using the multiband-accelerated EPI sequence (acceleration factor = 4, TR = 640 ms, TE = 30 ms, slices = 40, FOV = 192, voxel sizes = 3×3×3 mm).

The fMRI data were pre-processed using AFNI software^62^. The first 8 volumes were removed. For univariate analysis, the following preprocessing steps included slice timing, motion correction, aligning the functional image with anatomy, spatial normalization (MNI152 space), spatial smoothing with a 6-mm FWHM isotropic Gaussian kernel, and scaling each voxel time series to have a mean of 100. The fMRI data were not smoothed and scaled for MVPA and neural alignment analysis in the preprocessing steps.

### Univariate general linear model analysis

Single-subject multiple-regression modeling was performed using the AFNI program 3dDeconvolve. Four syllables under 3 signal-to-noise ratios and 6 regressors corresponding to motion parameters were entered into the analysis. TRs were censored if the motion derivatives exceeded 0.3. For each signal-to-noise ratio, the four syllables were grouped and contrasted against the baseline. Group-level analysis was performed using the AFNI program 3dMVM. To investigate how aging affects the neural response to audiovisual speech among musicians and non-musicians, 2 separate models were constructed with the same within-subject factor (signal-to-noise ratio) and different between-subject factors (ONM vs. YNM, OM vs. YNM). To investigate the musical training effect among older adults, one within-subject factor (signal-to-noise ratio) and one between-subject factor (subject group: OM vs. ONM) were put into the model. Multiple comparisons were corrected using 3dClustSim (“fixed” version) with real smoothness of data estimated by 3dFWHMx (acf method)^63^. Ten thousand Monte Carlo simulations were performed to get the cluster threshold (alpha = 0.05 family-wise-error corrected). Results were visualized onto a cortical inflated surface using SUMA with AFNI.

Conjunction analysis was performed to determine the regions that showed common and different aging effects between musicians and non-musicians. Corrected group analysis results masks (ONM vs. YNM, OM vs. YNM) were entered into the conjunction analysis, and the conjunction map was created by 3dcalc.

### Searchlight MVPA

An exploratory whole-brain searchlight MVPA with a 6-mm radius was conducted to decode the 4 syllables under each signal-to-noise ratio using the Decoding Toolbox^64^. The classifiers were trained using support vector machine algorithm with a linear kernel. The cost parameter C was set to 1. The input feature was univariate trial-wise β coefficients that were estimated using AFNI program 3dLSS which was recommended for performing MVPA in fast event-related designs^65^. Leave-one-run-out cross-validation was used to evaluate classification performance, which was measured by mean classification accuracy. Then, accuracy maps were smoothed with an 8-mm FWHM Gaussian kernel. The smoothed accuracy maps were entered into the group analysis using the AFNI program 3dMVM. The group analysis was the same as the procedure in the univariate analysis.

### Neural alignment-to-young analysis

The same single-subject multiple-regression modeling procedure as the univariate analysis was performed using functional data that were not smoothed and scaled. T-statistic maps of each syllable under 3 signal-to-noise ratios for each subject were used in the following neural alignment analysis.

Five significant clusters in the MVPA group analysis were employed as ROIs in the neural alignment analysis. The neural responses (t value) of each voxel within the ROIs to each syllable under 3 signal-to-noise ratios were extracted for each subject. Then, the neural responses of each voxel within the ROIs were averaged across all YNM, which yielded the mean activation pattern of young adults to each syllable under 3 signal-to-noise ratios. Then, for each older subject and each activation pattern within the ROI, we calculated the alignment-to-young measure, which is the spatial activation similarity between each older subject and young average when they are perceiving each audiovisual syllable, using Pearson correlation (see Fig. 3c). To simplify the analysis, 12 correlation coefficients (4 syllables * 3 signal-to-noise ratios) were averaged for each older subject. Such an inter-subject pattern correlation framework has been employed in previous studies related to memory and learning^36,66^. The alignment-to-young measure of OM and ONM were compared using the 2-sample t-test. FDR correction was performed to correct multiple comparisons.

### Correlation analysis

Correlation analyses were further conducted among BOLD responses, alignment-to-young measurement, and speech-in-noise performance in two older groups. ROIs that exhibited significant group differences in BOLD activity or alignment-to-young were selected. Since the neural responses of brain regions within the same network were highly correlated (Supplementary Fig. 2), to simplify the analysis we employed the principal component analysis with the varimax rotation to obtain the first principal component of each network that represented the correlated brain regions and was used in the correlation analysis.

Although bilateral auditory and visual areas were not within the same neural network, their activations were significantly correlated across subjects (see Supplementary Fig. 2). Therefore, they were projected onto 1 sensory component. The bilateral auditory and visual regions of OM were extracted from the significant clusters from the contrast between OM and YNM, and regions of ONM were extracted from the significant clusters from the contrast between ONM and YNM. The loadings of all 4 ROIs on the sensory component exceeded 0.65, which was moderate to high factor loading. BOLD responses in DMN regions including bilateral angular gyrus, medial prefrontal cortex, and ACC were projected onto 1 DMN component. The ACC responses of OM and ONM were also extracted among significant clusters that showed significant differences between each older group and YNM. The left angular gyrus responses of both groups were from the significant cluster of OM vs. ONM because this region was also significantly different between OM and YNM but not between ONM and YNM. The right angular gyrus and bilateral medial prefrontal cortex responses of both groups were from significant clusters of OM vs. YNM because those regions were not significant in the contrast between ONM and YNM. The loadings of all 4 ROIs on the DMN component exceeded 0.74. Responses of speech motor areas were not processed with principal component analysis because only 1 cluster covering the left IFGop and prCG that showed a significant difference between OM and ONM was included. Alignment-to-young measures in bilateral sensorimotor areas were projected onto 1 alignment component, with the loadings of both ROIs exceeding 0.91.

All principal components, speech motor BOLD response, and the mean audiovisual speech-in-noise accuracy (across conditions) were pairwise entered into the Bayesian Pearson correlation analysis. Bayesian Pearson correlation analysis was performed in R (R Core Team, 2017) with the package Correlation^67^. BF above 3 was considered moderate evidence for a significant correlation, and BF above 10 was considered strong evidence.

## Acknowledgments

The authors thank Yining Chen, Yiyang Wu for helping with data collection, and Xintong Jiang for helping with stimuli generation.

## Author contributions

L. Zhang acquired and analyzed the data. L. Zhang and Y. Du designed the experiment. L. Zhang, X. Wang, and Y. Du interpreted the results and wrote the manuscript.

## Data and materials availability

The data that support the findings of this study are available in https://osf.io/2wxhv/.

