## Supplementary materials for "Musical expertise enhances neural alignment-to-young in sensorimotor regions that predicts older adults’ audiovisual speech-in-noise perception"

**Effect of years of education on the behavioral performance and neural response in the audiovisual speech-in-noise perception task**

We found that years of education including both formal and informal education were significantly different between ONM and OM. Since all recruited older adults quitted the formal education at the same time (1960s-1970s) because of the historical reason in China, the self-reported years of education cannot represent the education level of the older subjects reliably.

Since years of education were highly correlated with the group variable between OM and ONM (point-biserial correlation r = 0.64), and were perfectly correlated with the group variable between older subjects and young subjects (ONM vs. YNM: $r_{pb}$= 1; OM vs. YNM: $r_{pb}$= 0.74), it is not suitable to put the years of education variable into the analyses as a control variable because of the collinearity problem.

To exclude the effect of years of education on our results, we performed supplementary analysis to investigate the effect of years of education on the behavioral performance, BOLD activation, and neural representation in older adults. We put the years of education in the mixed-designed ANOVA that analyzed the group effect of OM and ONM on the audiovisual speech perception performance. Years of education were not related to behavioral performance (*F*(1, 47) = 0.11, *p* = 0.738, $\eta_{p}^{2}=0.00$). Similarly, we added years of education in the group analysis of BOLD activation and MVPA classification accuracy between OM and ONM. No cluster showed significant effect of years of education even under a loose threshold and correction level ($p_{uncorrected}$ < 0.01, $p_{fwe}$ < 0.1). Therefore, years of education did not influence the behavioral performance and neural response of older adults in this study.
