## Supplementary Fig. 1 for "Musical expertise enhances neural alignment-to-young in sensorimotor regions that predicts older adults’ audiovisual speech-in-noise perception"

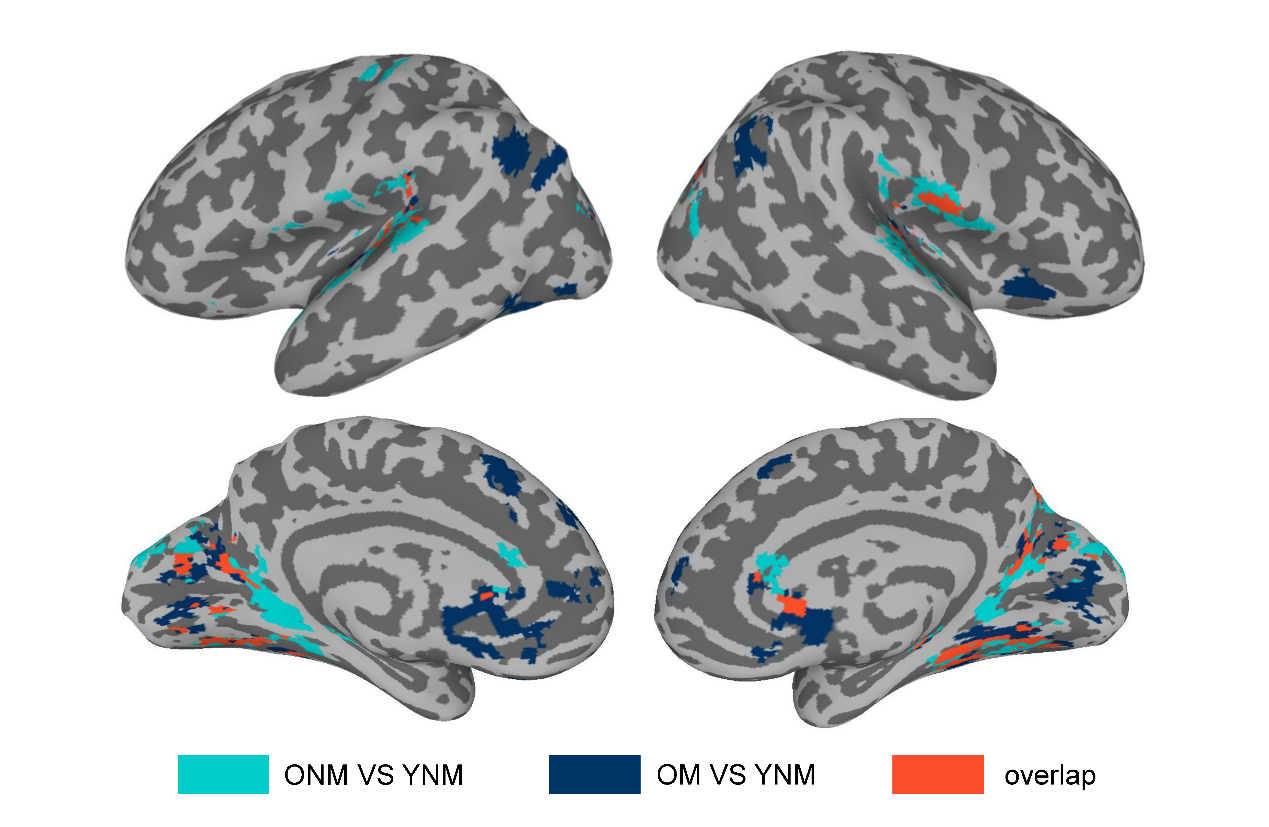

**Figure S1.** **Conjunction analysis of group comparison**. Brain regions in light blue show the specific significant regions of ONM vs. YNM. Brain regions in dark blue show the specific significant regions of OM vs. YNM. Brain regions in orange show the overlapping significant regions of ONM vs. YNM and OM vs. YNM. Clusters of fewer than 20 voxels were not shown here. OM: older musicians; ONM: older non-musicians; YNM: young non-musicians.
