## Supplementary Fig. 2 for "Musical expertise enhances neural alignment-to-young in sensorimotor regions that predicts older adults’ audiovisual speech-in-noise perception"

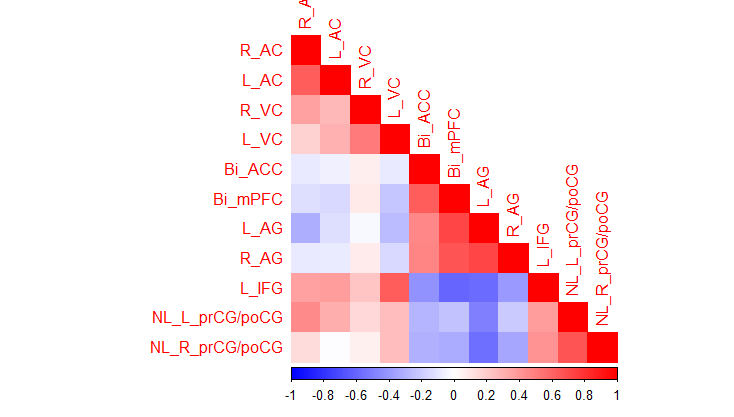

**Figure S2.** **Correlation matrix of the neural indices used in the Method section “Correlation analysis”**. The last two variables that start with NL were neural alignment-to-young measure, and the other measures were BOLD activation. Colors represent the Pearson correlation coefficient. R, right; L, left; Bi, bilateral. For abbreviations of brain regions, see Table S4.
