## Supplementary Table 1 for "Musical expertise enhances neural alignment-to-young in sensorimotor regions that predicts older adults’ audiovisual speech-in-noise perception"

**Table S1.** **Group analysis results of BOLD activity.** Brain regions showing significant group effect on BOLD activity during audiovisual speech-in-noise perception ($p_{fwe}$< 0.05). OM: older musicians; ONM: older non-musicians; YNM: young non-musicians. L, left; R, right; Bi, bilateral. For abbreviations of brain regions, see Table S4.

|  | **Brain Regions** | **Peak MNI coordinate** | | | **Peak t-value** | **No. of voxel** |
| --- | --- | --- | --- | --- | --- | --- |
|  |  | **x y z** | | |  |  |
| **OM > ONM** | L poCG/IPL/SPL | -41 | -50 | 58 | 5.43 | 208 |
|  | L SMG/STG | -59 | -23 | 10 | 4.89 | 172 |
|  | L PMC/IFGop | -53 | 4 | 37 | 4.04 | 71 |
| **OM < ONM** | L AG | -50 | -59 | 28 | -4.81 | 186 |
| **YNM > OM** | Bi ACC/OFC/CN/OC | -2 | 22 | 1 | 5.02 | 294 |
|  | L AG | -50 | -68 | 31 | 4.91 | 165 |
|  | L SFG/SFGmed | -20 | 55 | 31 | 4.93 | 151 |
|  | R AG/IPL | 58 | -59 | 37 | 4.54 | 108 |
|  | R ROL/HG | 58 | -8 | 13 | 5.09 | 95 |
|  | Bi SFGmed/ACC | -8 | 49 | 7 | 4.22 | 90 |
|  | L STG/ROL | -44 | -32 | 13 | 4.43 | 77 |
| **YNM < OM** | BI CALG/LING/CUN | 19 | -59 | 22 | -5.94 | 519 |
|  | R FFG/LING | 25 | -41 | -11 | -5.86 | 278 |
|  | L MOG | -29 | -83 | 28 | -5.05 | 117 |
|  | L ITG/IOG | -53 | -62 | -11 | -4.8 | 116 |
|  | R MOG | 34 | -80 | 28 | -5.53 | 94 |
|  | Bi SMA/SFGmed | -8 | 22 | 49 | -5.27 | 89 |
|  | R IFGorb/INS | 28 | 25 | -8 | -5.87 | 68 |
| **YNM > ONM** | R ROL/STG/HG | 61 | -8 | 10 | 8.6 | 338 |
|  | L STG/ROL/SMG | -59 | -32 | 13 | 5.46 | 200 |
|  | Bi ACC/R CN | 7 | 19 | -2 | 5.94 | 111 |
|  | L STG/TP | -53 | -8 | 1 | 5.12 | 86 |
|  | L prCG/poCG | -35 | -29 | 64 | 4.78 | 81 |
| **YNM < ONM** | Bi CUN/PCUN/CALG | 19 | -53 | 13 | -4.98 | 458 |
|  | L LING/FFG/CALG | -32 | -49 | -11 | -4.8 | 289 |
|  | R LING/FFG/PHG | 28 | -38 | -11 | -8.04 | 221 |
|  | R MOG | 34 | -83 | 28 | -5.56 | 141 |
|  | L MOG | -26 | -87 | 30 | -4.57 | 98 |
