## Supplementary Table 2 for "Musical expertise enhances neural alignment-to-young in sensorimotor regions that predicts older adults’ audiovisual speech-in-noise perception"

**Table S2.** **Conjunction analysis results of group effect on BOLD activity.** Overlapping: overlapping regions between OM vs. YNM and ONM vs. YNM. OM specific: specific regions that OM showed significant difference of BOLD activity compared to YNM. ONM specific: specific regions that ONM showed significant difference of BOLD activity compared to YNM. Coordinates for the center of each cluster were reported. OM: older musicians; ONM: older non-musicians; YNM: young non-musicians. L, left; R, right; Bi, bilateral. For abbreviations of brain regions, see Table S4.

|  | **Brain Regions** | **Center MNI coordinate** | | | **No. of voxel** |
| --- | --- | --- | --- | --- | --- |
|  |  | **x** | **y** | **z** |  |
| **Overlapping** | R FFG/LING | 25 | -50 | -11 | 127 |
|  | R ROL/HG | 49 | -14 | 10 | 78 |
|  | L FFG/LING | -26 | -62 | -8 | 76 |
|  | R MOG | 34 | -80 | 25 | 74 |
|  | R CUN/PCUN | 13 | -65 | 22 | 71 |
|  | L MOG | -29 | -83 | 25 | 59 |
|  | R CN/OC/Bi ACC | 7 | 22 | -2 | 54 |
|  | L STG | -47 | -32 | 13 | 52 |
|  | L CALG | -20 | -65 | 13 | 27 |
| **OM specific** | L ACC/OFC | -2 | 28 | -8 | 222 |
|  | Bi CALG/LING | -2 | -74 | 10 | 203 |
|  | L AG | -50 | -65 | 31 | 165 |
|  | L SFG/OFC | -14 | 52 | 34 | 151 |
|  | R FFG/LING | 22 | -53 | -14 | 138 |
|  | L ITG/IOG | -50 | -65 | -11 | 116 |
|  | R AG | 52 | -62 | 37 | 108 |
|  | Bi SFGmed/ACC | -2 | 55 | 7 | 90 |
|  | Bi SMA/SFGmed | -5 | 22 | 49 | 89 |
|  | R IFGorb/INS | 31 | 25 | -8 | 68 |
|  | R CUN/PCUN | 16 | -65 | 22 | 49 |
|  | L MOG | -32 | -83 | 31 | 47 |
| **ONM specific** | Bi CUN/SOG | 1 | -86 | 22 | 269 |
|  | R ROL/STG | 55 | -14 | 7 | 257 |
|  | L LING/CALG | -17 | -50 | -2 | 138 |
|  | L STG/SMG | -56 | -23 | 13 | 134 |
|  | R LING/FFG/ | 25 | -50 | -5 | 84 |
|  | L poCG/prCG | -38 | -26 | 58 | 81 |
|  | L STG/TP | -50 | 4 | -5 | 80 |
|  | R MOG | 40 | -80 | 22 | 60 |
|  | R CALG/PCUN | 19 | -59 | 13 | 49 |
|  | Bi ACC | 1 | 31 | 16 | 41 |
|  | R CUN/SOG | 16 | -84 | 48 | 31 |
