## Supplementary Table 3 for "Musical expertise enhances neural alignment-to-young in sensorimotor regions that predicts older adults’ audiovisual speech-in-noise perception"

**Table S3.** **MVPA group analysis results.** Brain regions showing significant group effect on the classification accuracy ($p_{\mathrm{fwe}}$< 0.05). OM: older musicians; ONM: older non-musicians; YNM: young non-musicians. L, left; R, right. For abbreviations of brain regions, see Table S4.

|  | **Brain Regions** | **Peak MNI coordinate** | | | **Peak t-value** | **No. of voxel** |
| --- | --- | --- | --- | --- | --- | --- |
|  |  | x | y | z |  |  |
| **OM > ONM** | L LING/MOG/IOG | -30 | -86 | -12 | 5.1 | 470 |
|  | L prCG/IFGop | -33 | 2 | 26 | 4.46 | 367 |
| **YNM > ONM** | L prCG/poCG/SMA | -33 | -22 | 64 | 4.85 | 1167 |
|  | L LING/MOG/IOG | -24 | -89 | -12 | 5.15 | 468 |
|  | R prCG/poCG | 27 | -15 | 61 | 4.65 | 434 |
