## Supplementary Table 4 for "Musical expertise enhances neural alignment-to-young in sensorimotor regions that predicts older adults’ audiovisual speech-in-noise perception"

**Table S4. Names and abbreviations of the brain regions used in the article and supplementary materials.**

| Brain regions | Abbreviation |
| --- | --- |
| angular gyrus | AG |
| anterior cingulate cortex | ACC |
| calcarine gyrus | CALG |
| caudate nucleus | CN |
| cuneus | CUN |
| fusiform gyrus | FFG |
| heschl's gyrus | HG |
| inferior frontal gyrus, opercular part | IFGop |
| inferior frontal gyrus, orbital part | IFGorb |
| inferior occipital gyrus | IOG |
| inferior parietal lobule | IPL |
| inferior temporal gyrus | ITG |
| insula | INS |
| lingual gyrus | LING |
| middle occipital gyrus | MOG |
| olfactory cortex | OC |
| orbitofrontal cortex | OFC |
| postcentral gyrus | poCG |
| precentral gyrus | prCG |
| precuneus | PCUN |
| premotor cortex | PMC |
| rolandic operculum | ROL |
| superior frontal gyrus, medial | SFGmed |
| superior occipital gyrus | SOG |
| superior temporal gyrus | STG |
| supplementary motor area | SMA |
| supramarginal gyrus | SMG |
| temporal lobe | TP |
